# Increased Genetic Protection Against Alzheimer’s Disease in Centenarians

**DOI:** 10.1101/2025.06.02.657442

**Authors:** Harold Bae, Zeyuan Song, Amanat Ali, Takashi Sasaki, Niccolò Tesi, Hannah Lords, Anastasia Leshchyk, Yukiko Abe, Nobuyoshi Hirose, Yasumichi Arai, Nir Barzilai, Erica F. Weiss, Marc Hulsman, Sven van der Lee, Natasja M. van Schoor, Martijn Huisman, Yolande Pijnenburg, Wiesje van der Flier, Marcel Reinders, Henne Holstege, Sofiya Milman, Thomas T Perls, Stacy L Andersen, Paola Sebastiani

## Abstract

We constructed a polygenic protective score specific to Alzheimer’s disease (AD PPS) based on the current literature among the participants enrolled in five studies of healthy aging and extreme longevity in the US, Europe, and Asia. This AD PPS did not include variants on Apolipoprotein E (APOE) gene. Comparisons of AD PPS in different data sets of healthy agers and centenarians showed that centenarians have stronger genetic protection against AD compared to individuals without familial longevity. The current study also shows evidence that this genetic protection increases with increasingly older ages in centenarians (centenarians who died before reaching age 105 years, semi-supercentenarians who reached age 105 to 109 years, and supercentenarians who reached age 110 years and older). However, the genetic protection was of modest size: the average increase in AD PPS was approximately one additional protective allele per 5 years of gained lifetime. Additionally, we show that the higher AD PPS was associated with better cognitive function and decreased mortality. Taken together, this analysis suggests that individuals who achieve the most extreme ages, on average, have the greatest protection against AD. This finding is robust to different genetic backgrounds with important implications for universal applicability of therapeutics that target this AD PPS.

## INTRODUCTION

Alzheimer’s disease (AD) is the most common cause of dementia with a high heritability ranging from 60% to 80% (1,2). Centenarians demonstrate exceptional health spans by delaying many age-related diseases (3–5). Paradoxically, centenarians also have an increased risk of AD because of their exceptional age, but are often able to delay or avoid AD and related dementias (6). Therefore, they may serve as an important model to study the mechanisms of resilience or resistance to AD and to identify genetic factors for neuroprotection from AD.

While the effect of the apolipoprotein E (*APOE*) gene on AD is well-characterized, the search for additional reliable genetic factors for AD has been ongoing. A recent GWAS analysis (7) identified a total of 83 genetic variants associated with AD using 111,326 clinically diagnosed/’proxy’ AD cases and 677,663 controls of White/European ancestry. In this list of genetic variants, 44 were novel loci at the time of publication. Given that individual single-nucleotide polymorphisms (SNPs) typically have a limited impact on disease risk, polygenic risk scores that aggregate the effect of multiple genetic loci have been developed for various human diseases and phenotypes (8). Recently, using the polygenic risk score approach with the above list of genetic variants, Tesi et al. showed that cognitively healthy centenarians are enriched with the protective genetic alleles associated with AD compared to AD cases and middle-aged healthy controls (9).

In the present study, we construct the AD-specific polygenic protective score (AD PPS) based on the list of 83 genetic variants that does not include the *APOE* variants in participants of the New England Centenarian Study (NECS) (10). We compared the distributions of the AD PPS between the centenarians, offspring of centenarians, and study controls. Furthermore, the exceptional ages of the NECS participants allowed us to compare the distributions in four age categories--nonagenarians, centenarians who died before reaching age 105 years, semi-supercentenarians who reached age 105 to 109 years, and supercentenarians who reached age 110 years and older. We examined the association between the AD PPS and performance on cognitive screeners (i.e., Blessed Information-Memory-Concentration [BIMC] test and modified Telephone Interview for Cognitive Status [TICS]) and mortality. The BIMC is a standardized cognitive screening test that assesses memory, working memory, and orientation, has a high completion rate in very old individuals(11,12), and was validated against neuropathology(11). The TICS is a cognitive screener that includes items to assess orientation, episodic memory, language, and working memory and has high sensitivity and specificity for AD (13). Both the BIMC and TICS are reliable instruments for determining cognitive impairment over the phone.

We replicated the analysis using data from the Long Life Family Study (LLFS) (14), Longevity Genes Project (LGP) (15) / LonGenity (16) Study cohorts, the Tokyo Centenarian Study (TCS) and the Japan Semisupercentenarian Study (JSS), and 100-plus Study by comparing the AD PPS between the centenarians, offspring of centenarians, and study controls and in different age groups. In the LLFS and LGP/LonGenity, we also examined the association between AD PPS and various neuropsychological tests and mortality.

## METHODS AND RESULTS

### Study Populations and Genetic Data

#### The New England Centerarian Study (NECS), The Long Life Family Study (LLFS), and LGP/Longenity Study

NECS is a study of extreme human longevity that includes centenarians, their long-lived siblings, offspring, offspring spouses and additional unrelated controls. Since the inception of the study in 1994, it has enrolled more than 2,500 centenarians including 600 semi-supercentenarians (ages 105-109) and 200 supercentenarians (ages 110+ years), and some of their siblings, about 500 offspring and about 350 controls from North America (10).

LLFS is a family-based study of healthy aging and longevity that recruited nearly 5000 individuals from 583 multigenerational families selected for familial longevity in the United States and Denmark. Selection criteria for participants have been described in detail elsewhere (14,17,18). Participants from the Denmark site were excluded from the current analysis due to the data restriction agreement.

The participants of LGP and LonGenity studies were of Ashkenazi Jewish descent as described elsewhere (15,19). The LGP established has recruited Ashkenazi Jewish centenarians (age 95 and older) who are in general good health at age 95, offspring of centenarians, and spouses of offspring since 1998. The LonGenity study has been recruiting community dwelling Ashkenazi Jewish seniors aged 65 or older in the United States since 2008, including Offspring of Parents with Exceptional Longevity (OPEL), defined by having at least one parent who lived to age 95 or older.

Genome-wide genotype data in the NECS, LLFS, LGP/LonGenity were imputed to the HRC panel (version r1.1 2016) of 64,940 haplotypes with 39,635,008 sites using the Michigan Imputation Server (20). Previous genetic analyses involving these three cohorts were described in (21–23). Participants in all these studies provided informed consents.

#### The Japanese Centenarian Study: TCS / JSS

For the Japanese centenarian group, we used data from two prospective cohort studies of the oldest individuals in Japan, the Tokyo Centenarian Study (TCS) and the Japan Semisupercentenarian Study (JSS) (24,25). The cut-off date for the data collected from the Cent group was May 31, 2022. Among the 967 centenarians from whom gDNA was obtained, 964 centenarians were included, with data from three centenarians who had mismatched sex information or were assigned to close relatives (PI HAT of 0.1875 or higher, which represents the half-way point between 2nd- and 3rd-degree relatives) excluded; this resulted in a study population comprising 144 men and 820 women (female-to-male ratio: 0.851) with a median age of 106.0 years [interquartile range (IQR): 103.9-107.1]. Among these individuals, there were 915 deaths (death rate: 0.969), with a median age at death of 107.5 years [IQR: 105.7-109.2].

#### 100-plus Study / ADC / LASA

Cognitively healthy centenarians were recruited from the 100-plus Study, which includes Dutch-speaking individuals who can provide evidence for being aged 100 years or older, and self-report to be cognitively healthy (26). Genetic variants were determined as previously described in (9) following established quality control and imputation methods (9,20,27,28). Imputation was performed using the TOPMED imputation panel ((29), see Supplementary Methods). Before analyses, we excluded individuals with a family relation (identity-by-descent >0.2, except for the centenarians’ offspring which were all included), and only individuals of European ancestry (based on 1000Genome clustering) were kept.

In the NECS, LLFS, and LGP and LonGenity, we defined offspring of a centenarian as any participant with at least one parent who survived to age 100 or older. In the 100-plus study, offspring were offspring of the cognitively healthy centenarians. In the NECS, study controls were referent participants in the NECS who were spouses of centenarian offspring or children of individuals who died at approximately age 73 years and matched the life expectancy of their birth cohort (30). In the LLFS, study controls were the spouses of the family members. In the LGP and LonGenity, study controls were individuals without a parental history of longevity (neither parent survived beyond 95□years of age). In the 100-plus study, healthy controls were recruited from different cohorts (see Supplementary Methods) (26,31–34).

### Construction of AD PPS

We initially considered 83 genetic variants (Supplementary Table 5 in (7)) identified in a large-scale GWAS meta-analysis with 111,326 clinically diagnosed/’proxy’ AD cases and 677,663 controls that are of White/European ancestry (7). However, adequate proxy SNPs (r^2^ > 0.8) for nine variants could not be found, leading to a total of 74 genetic variants in the NECS, LLFS, and 100-plus Study, 59 variants in the LGP and LonGenity, and 60 variants in the TCS/JSS. We constructed the AD PPS by adding up the protective alleles (i.e. alleles that decreases the odds of having AD). Of note, the *APOE* variants were excluded from this analysis. We chose the unweighted summation approach as the effect sizes may not be transferable given the non-homogenous ethnic compositions of the included study cohorts (35).

### Measures: Cognitive Screeners and Neuropsychological Tests

In the NECS, there were two cognitive screeners (BIMC and TICS) available for analysis. For the BIMC test, there were 483 centenarians available for analysis. Among these participants, 233 had baseline measurements only, 133 had two measurements, and 117 had three or more repeated measurements. For the TICS test, we analyzed 637 offspring and controls in the NECS, and 534 participants had three or more repeated measurements.

In the LLFS, there were 29 outcomes of neuropsychological tests and cognitive screeners. The LLFS has administered comprehensive neuropsychological assessments three times. At enrollment (2006—2009), LLFS participants completed eight of the ten tests in the National Alzheimer’s Coordinating Center (NACC) Uniform Data Set (36), including the Mini Mental State Examination (MMSE), digit spans, logical memory, category fluency, and the Digit Symbol Substitution Test (DSST). LLFS completed a second administration of the same battery of tests and the protocol expanded to also include the Hopkins Verbal Learning Test – Revised, Clock Dawing Test (CDT), Trail Making Test (TMT), and phonemic fluency and added the use of a digital pen to record performance on all graphomotor tasks (i.e., (MMSE, CDT, DSST, and TMT).

We developed algorithms to differentiate test completion time into time used to think about the test answers (think time) and time used to write the answers (ink time) for both the digitally administered DSST (37) and Trail Making test (38). For the current study, the outcomes included the DSST, Digit Span score (forward and backward), Logical Memory score (immediate and delayed recall), and animal fluency score at visit 1 and visit 2 as well as the difference between the two measurements at both visits for each test. Additionally, cognitive trajectories based on the TICS scores and various measures of think time and ink time were included.

In the LonGenity study, neuropsychological tests included logical memory immediate recall, semantic fluency, digit span forward and backward, and DSST.

### Statistical Analysis

As a validation analysis, we compared the calculated AD PPS to the list of 54 polygenic risk scores for common diseases computed in the NECS and LLFS participants by our group (39). The analysis was adjusted for sex, top 4 principal components, and family relatedness in linear mixed effects models. Then, two main comparison analyses were performed. First, the distributions of the AD PPS were compared between the centenarians, offspring of centenarians, and controls in the NECS, LLFS, LGP/LonGenity, and 100-plus Study. Next, to examine whether the AD PPS differed as a function of increasing age, the distributions of the AD PPS were compared in four age categories—nonagenarians, centenarians (ages 100-104), semi-supercentenarians (ages 105-109), and supercentenarians (age 110+). The models adjusted for sex and top 4 principal components (where necessary). In the NECS and LLFS, the generalized estimation equations (GEE) models treating each family as a cluster were used to account for family relatedness. In the TCS/JSS, the distributions of the AD PPS were compared in three age categories—centenarians (ages 100-104), semi-supercentenarians (ages 105-109), and supercentenarians (age 110+).

In the NECS, we correlated the AD PPS on two measures of cognitive screeners (BIMC and TICS) by adjusting for age at the time of the test administration, sex, years of education, and APOE alleles. For the longitudinal, repeated measurements of BIMC and TICS, we used a random intercept model and also included the interaction between the AD PPS and age at the time of administration to assess the rate of change. In the LLFS, we analyzed the effect of the AD PPS on 29 neuropsychological and cognitive test scores using regression adjusted for the interaction between age at consent and generation, sex, education, top 4 principal components, APOE alleles, and a random intercept per family. The generation variable indicates whether a participant belongs to the proband or offspring generation. In the LonGenity, the effect of the AD PPS on 5 neuropsychological tests were assessed using regression adjusted for age, sex, and education.

In the NECS and LLFS, we also examined the association between the AD PPS and mortality. We conducted a Cox proportional hazard regression analysis of age at death (or censored at age of last contact) with the AD PPS as the main predictor, adjusting for sex and top 4 principal components. A random effect term per family (frailty model) was included to account for family correlations. We assessed the proportional hazards assumptions by testing for independence between the scaled Schoenfeld residuals and time, accompanied by visual inspection of scaled Schoenfeld residuals by time. In the NECS, we determined that participants’s sex may have violated the proportional hazards assumption, and thus we fit a stratified Cox model.

The LLFS underwent a comprehensive dementia review on a smaller subset of participants (n=1782) based on a thorough evaluation of cognitive conditions. We used the probable Alzheimer’s disease by NINCDS-ADRDA criteria (40). In a logistic regression model, we test whether the AD PPS is predictive of the AD diagnosis adjusting for the interaction between age at last contact and generation, sex, and top 4 principal components with a random intercept to account for family relatedness. In this analysis, there were 250 AD cases and 1532 non-AD controls.

## RESULTS

**Table 1** sumamrizes the cohorts included in this study. In the NECS, there were 1255 centenarians, 478 offspring of centenarians, and 306 controls. There were 437 nonagenarians, 610 centenarians who died before reaching age 105, 516 semi-supercentenarians who reached age 105 to 109, and 134 supercentenarians who reached age 110 years and older. In the LLFS, there were 379 centenarians, 432 offspring of centenarians, and 505 controls. There were 976 nonagenarians, 376 centenarians who died before reaching age 105, 69 semi-supercentenarians, and 5 supercentenarians. In the LGP and LonGenity, there were 299 centenarians, 243 offspring of centenarians, and 509 controls. There were 260 nonagenarians, 198 centenarians who died before reaching age 105, 93 semi-supercentenarians, and 8 supercentenarians. In the TCS/JSS, there were 171 centenarians who died before reaching age 105, 577 semi-supercentenarians who reached age 105 to 109, and 161 supercentenarians who reached age 110 years and older. In the 100-plus Study, there were 3,165 healthy controls, 347 cognitively healthy centenarians, and 193 centenarians’ offspring for the analyses.

**Table 1.**
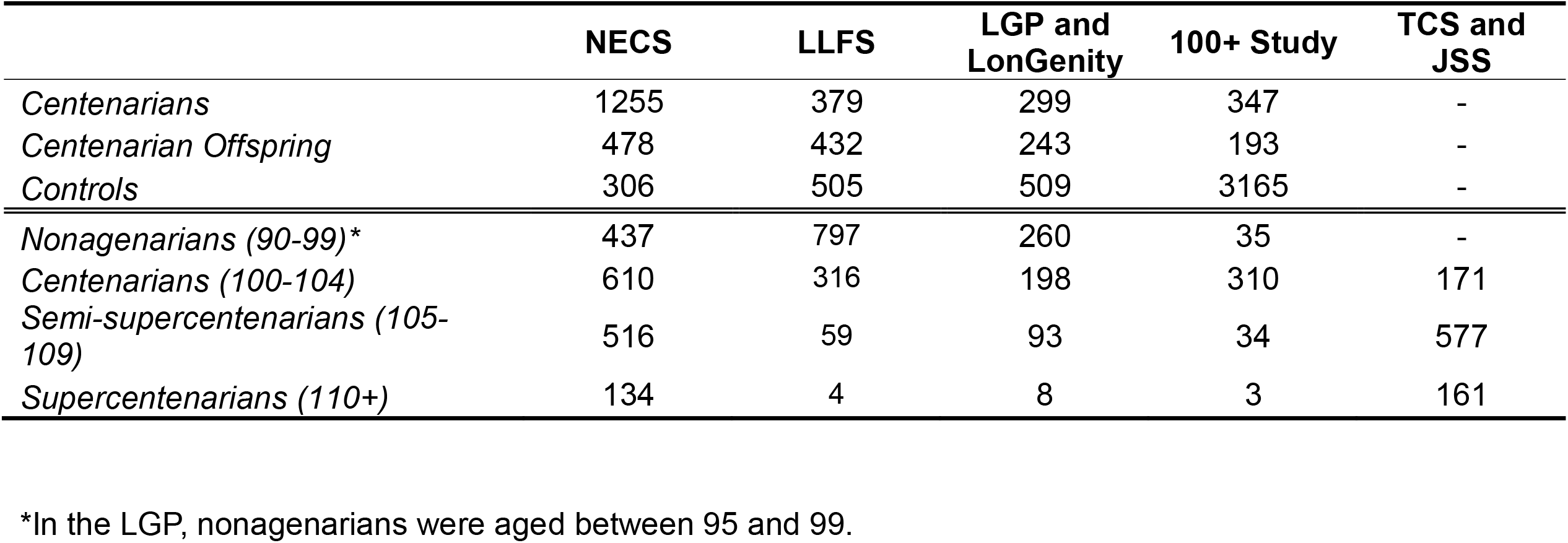
Characteristics of Study Cohorts.

In these cohorts, we generated the AD PPS as described in the methods. We used 74 SNPs in the NECS, LLFS and 100-plus Study, 60 SNPs in the TCS, and 59 SNPs in the LGP and LonGenity. The ranges of the calculated AD PPS differ by cohorts because different numbers of SNPs were used to calculate the scores.

### Validation of AD PPS in the NECS and LLFS

When the AD PPS was compared to the list of 54 polygenic risk scores for common diseases in the NECS and LLFS participants (39), we observed that, in both the NECS and LLFS, the current AD PPS had the strongest correlation with the PRS that is specific to AD in (39) with p=1.79×10^−55^ in the NECS and p=4.08×10^−70^ in the LLFS. The list of top 10 PRS’s associated with the current AD PPS in the NECS and LLFS are shown in Supplementary Table S1.

### Centenarians exhibit stronger genetic protections for AD than younger individuals

Table 2 summarizes the distribution of the AD PPS in the four cohorts included in this study. In the NECS, we observed that centenarians (mean AD PPS: 78.3) and offspring (mean AD PPS: 77.9) had significantly higher AD PPS than controls (mean AD PPS: 76.8) (p=3×10^−5^ and p=0.0081, respectively). The difference between centenarians and offspring AD PPS was not significant (p=0.145). In the LLFS, centenarians (mean AD PPS: 77.6) had significantly higher AD PPS than controls (mean AD PPS: 77.1) (p=0.04). The difference between offspring (mean AD PPS: 77.1) and controls was not significant (p=0.69) and the difference between centenarians and offspring was marginally significant (p=0.09). In the LGP and LonGenity, centenarians (mean AD PPS: 62.2) had higher AD PPS than the controls (mean AD PPS: 61.5), but the result was marginally significant (p=0.08). Other comparisons were not statistically significant in the LGP and LonGenity. Finally, in the 100-plus Study, centenarians (mean AD PPS: 78.3) had higher AD PPS than controls (mean AD PPS: 77.3) (p=0.000081) and offspring (mean AD PPS: 77.3) (p=0.02), but the difference between offspring and controls did not reach nominal significance.

**Table 2.** Comparison of AD PPS among centenarians, offspring of centenarians, and controls in different study cohorts.

|  | **NECS** |  | **LLFS** |  | **LGP and LonGenity** |  | **100+ Study** |  |
| --- | --- | --- | --- | --- | --- | --- | --- | --- |
|  | N | Mean AD PPS | N | Mean AD PPS | N | Mean AD PPS | N | Mean AD PPS |
| Controls | 306 | 76.8 | 505 | 77.1 | 509 | 61.5 | 3165 | 77.3 |
| Offspring | 478 | 77.9 | 432 | 77.1 | 243 | 62.1 | 193 | 77.3 |
| Centenarians | 1255 | 78.3 | 379 | 77.6 | 299 | 62.2 | 347 | 78.3 |

### Genetic protections for AD increases with extreme old ages

Table 3 summarizes the distributions of the AD PPS when the oldest participants (ages >90 years) were stratified in four age categories. In the NECS participants, we observed an increasing trend in the AD PPS with increasing age; the mean AD PPS in the nonagenarians, centenarians, semi-supercentenarians, and supercentenarians were 77.89, 77.97, 78.58, and 79.14, respectively (see Figure 1). While the AD PPS of centenarians and nonagenarians were not significantly different (p=0.85), the difference of AD PPS between semi-supercentenarians and centenarians was statistically significant (p=0.047). Similarly, the difference in AD PPS between super-centenarians and centenarians was statistically significant (p=0.023).

**Table 3.** Comparison of AD PPS in different age groups in different study cohorts.

|  | NECS |  | LLFS |  | LGP and LonGenity |  | TCS/JSS |  | 100+ Study |  |
| --- | --- | --- | --- | --- | --- | --- | --- | --- | --- | --- |
|  | N | Mean AD PPS | N | Mean AD PPS | N | Mean AD PPS | N | Mean AD PPS | N | Mean AD PPS |
| Nonagenarians (90-99)* | 437 | 77.89 | 797 | 77.52 | 260 | 61.8 | NA | NA | 35 | 78.2 |
| Centenarians (100-104) | 610 | 77.97 | 316 | 77.4 | 198 | 61.9 | 171 | 53.9 | 310 | 78.4 |
| Semi-supercentenarians (105-109) | 516 | 78.58 | 59 | 78.55 | 93 | 62.3 | 577 | 54.3 | 34 | 77.2 |
| Supercentenarians (110+) | 134 | 79.14 | 4 | 79.65 | 8 | 62.5 | 161 | 54.2 | 3 | 79.2 |

\*In the LGP, nonagenarians were aged between 95 and 99.

**Figure 1.**
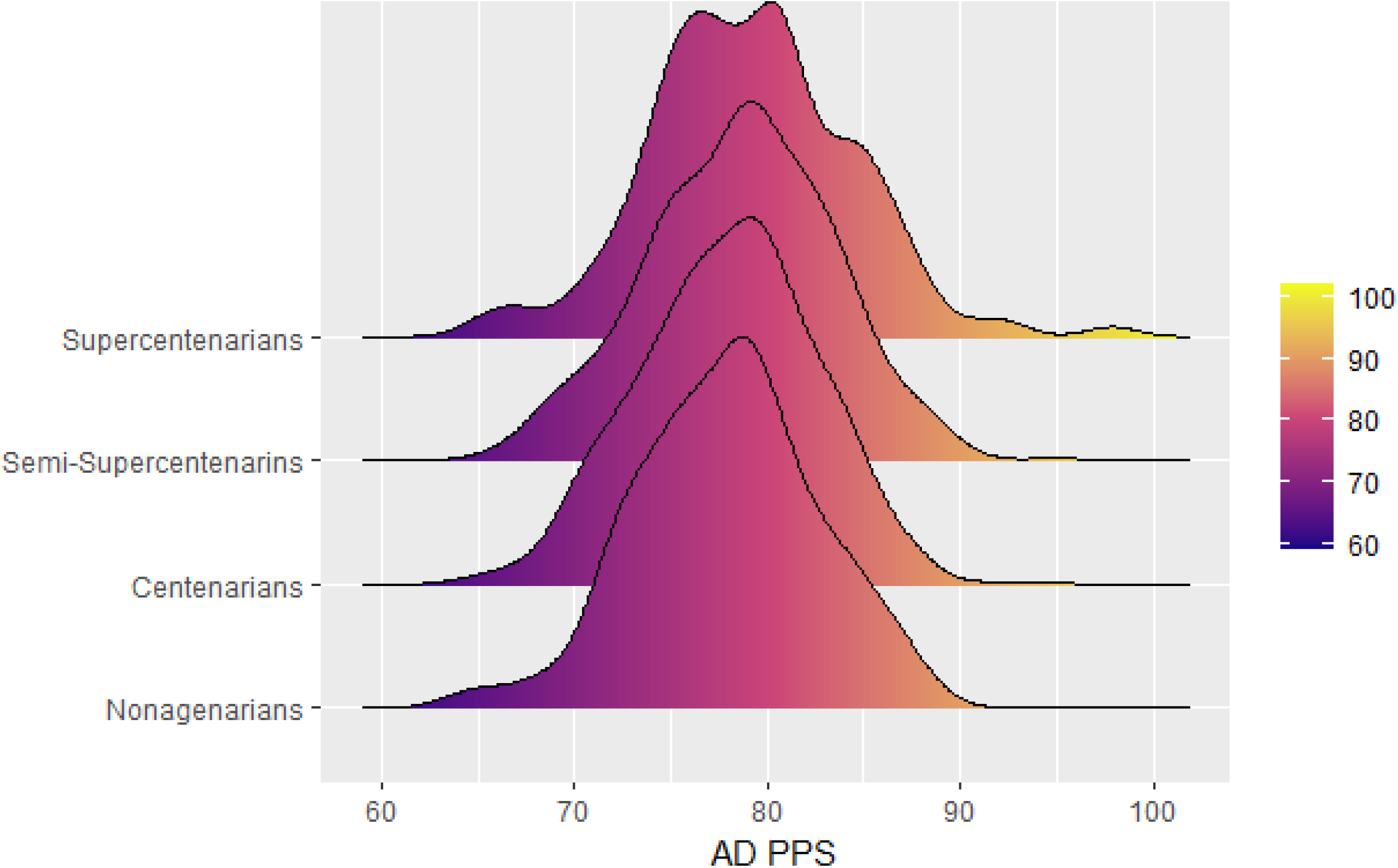
Comparison of the distributions of AD PPS nonagenarians (age 90-99), centenarians (100-104), semi-supercentenarians (age 105-109), and supercentenarians (age 110+) in the NECS.

Similar general patterns were observed with the LLFS, LGP, and TCS/JSS, although not every comparison yielded statistically significant results. In the LLFS, the mean AD PPS in the nonagenarians, centenarians, semi-supercentenarians, and supercentenarians were 77.52, 77.40, 78.55, and 79.65, respectively. There was no significant difference in the AD PPS between nonagenarians and centenarians, but semi-supercentenarians had a marginally significantly higher AD PPS than nonagenarians (p=0.07) and centenarians (p=0.08). Given the low sample size in the supercentenarians in the LLFS (n=4), we did not make any formal comparisons with this group. In the LGP, we again observed an increasing trend in the AD PPS with increasing age. However, none of the comparisons yielded statistically significant results. In the TCS/JSS, semi-supercentenarians had higher AD PPS than centenarians (p=0.004), but the difference between semi-supercentenarians and supercentenarians was not statistically significant. The limited number of centenarians older than 104 years in the 100-plus study did not produce any significant comparison.

### Higher AD PPS correlated with better cognitive function in centenarians

In NECS, we detected a significant association between the AD PPS and BIMC scores at enrollment in centenarians (beta=0.22, p=0.0026) but no association between AD PPS and rate of decline of BIMC scores. We did not detect a significant association between AD PPS and TICS scores in centenarians’ offspring and controls, although the TICS score increased with increasing AD PPS (beta=0.047, p=0.20). In the repeated measurement analysis, we did not find any evidence of association between AD PPS and rate of decline in the TICS scores. Results from the BIMC and TICS analyses in the NECS are summarized in Supplementary Table S2.

When we correlated the AD PPS with 29 neuropsychological test scores in the LLFS, the AD PPS was significantly associated with cognitive trajectories based on the TICS scores (beta=0.0014, p=0.013) and the difference in the logical memory delayed recall between visit 1 and 2 (beta=0.0396, p=0.027), but was not associated with their baseline scores. In both results, the positive beta’s indicate that individuals with higher AD PPS had slower decline in cognition. Full results of all 29 outcomes are shown in Supplementary Table S3. When AD PPS was correlated with the AD diagnosis in the LLFS, we observed that higher AD PPS reduces the odds of being diagnosed with AD (odds ratio=0.97), but the result was not statistically significant (p=0.17). In the LonGenity, none of the neuropsychological tests was associated with AD PPS.

### Increasing genetic protection from AD correlates with delayed mortality

We used Cox proportional hazard regression to correlate the AD PPS with mortality risk. This analysis showed that higher AD PPS was associated with lower hazard ratio (HR) for mortality (HR = 0.976, p=1.40×10^−6^) in the NECS. Separate analyses of the centenarians, offspring, and controls revealed that this significant association was driven by the centenarians. The HR’s for the three models were HR=0.979 (p=0.00014), HR=0.989 (p=0.5), and HR=0.990 (p=0.58), respectively for centenarians, offspring, and controls. The survival advantage for those with higher AD PPS is illustrated in Figure 2A.

**Figure 2.**
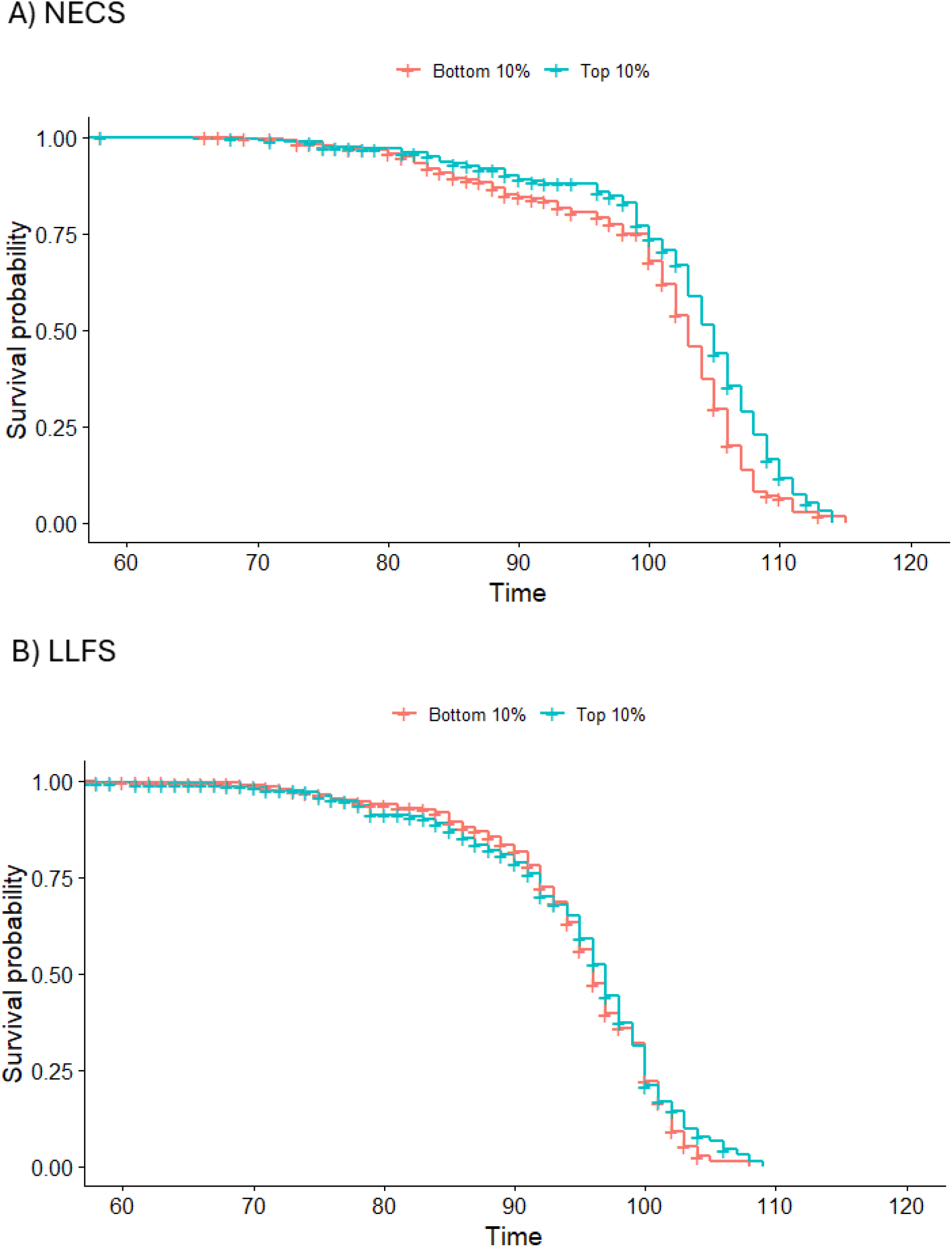
The Kaplan Meier curves comparing the top and bottom deciles of the AD PPS in the A) NECS and B) LLFS. In the NECS, there is a clear survival advantage among those in the top decile of the AD PPS compared to the participants in the bottom decile. In the LLFS, the survival advantage is more pronounced in participants who are 100 years or more.

This results replicated in the LLFS: a 1 point increase in the AD PPS was associated with HR=0.988 (p=0.018) in the overall sample. Figure 2B shows the Kaplan-Meier curves comparing the top and bottom deciles of the AD PPS in the LLFS participants. Note that the thresholds of the top and bottom deciles derived from the NECS were applied to the LLFS data.

## DISCUSSION

We constructed a polygenic protective score specific to AD based on the current literature (7) among the participants enrolled in studies of healthy aging and extreme longevity (10,15–17,22). We then examined the distributions of this AD PPS in different data sets and in different age groups and further correlated with various measures of cognitive function and mortality. The present study shows that there is stronger genetic protection against AD in centenarians compared to younger individuals and this result reproduced consistently in four additional cohorts with different study designs and genetic backgrounds. The AD PPS also correlated with cognitive decline and mortality in NECS and LLFS centenarians. In addition, our analysis showed that this genetic protection increases with increasingly older ages of centenarians, although the average difference was approximately one additional protective allele per 5 years of gained lifetime.

Previous work (9) showed that that cognitively healthy centenarians are enriched of genetic variants that protect from AD. Among the most replicated finding is an enrichment of the allele e2 of APOE in centenarians (41), and evidence that homozygosity for the e2 allele of APOE delays the onset of cognitive impairment (42). Besides APOE alleles, evidence of specific genetic protection from AD is sparse and points to a variety of possible biological pathways (22). The first article to show that cognitively intact centenarians are enriched for multiple genetic variants that protect from AD was published by the 100-plus Study in 2024 (9). Our work expands this result in two important directions. By including independent cohorts from the US, Europe, and Asia, our work shows that this genetic protection is robust to different genetic backgrounds. This finding is important for determining universal applicability of therapeutics that target this AD PPS.

The second important and novel finding of our work is evidence that the genetic protection becomes stronger with the older ages attained by centenarians. We showed in (4) that many centenarians delay the onset of morbidity and disability to the end of their lives and the gain of healthy longevity years increases with the older attained ages. In particular, supercentenarians who lived beyond 110 years appear to compress morbidity to the very end of their lives. The results in this paper are consistent with those findings and suggest that genetic protection may be one of the factors that enable some individuals to live to extreme old ages.

Our analysis also highlights a few surprising facts. Although statistically significant, the increase in the AD PPS by age groups was very small. For example, in the NECS the average AD PPS in nonagenarians and centenarians who died before reaching the age of 105 years was approximately 78, it increased to approximately 78.5 in semi-super centenarians and to 79.2 in supercentenarians. This trend was consistent in the TCS/JSS, and the LGP although the range of the AD PPS was different because the number of SNPs available to generate this score was smaller. These differences in AD PPS suggest that supercentenarians, on average, may carry just one additional protective allele compared to referent individuals. It is surprising that such a small difference may have a substantial effect of compression of morbidity and risk for AD. Our conjecture is that the AD PPS is simply one component of the genetic protection from AD and centenarians’ genomes may harbor additional rare genetic variants that are not found in the general population. The LLFS provided some evidence to support this hypothesis, since the study identified a cluster of rare variants in chromosome 3 that associated with significantly higher DSST scores (43). In addition, LLFS recently identified a new gene associated with AD risk (44). LLFS investigators also showed that a cluster of study participants who exhibited exceptional performance across a variety of neuropsychological tests were enriched for genetic variants that protect from AD and dementia (45). These results suggest that the catalogue of robust genetic variants associated with AD is not complete, although comprehensive, and more work is needed to identify the variety of genetic protections that could provide new avenues of therapeutics.

An additional interesting finding was that a large portion of centenarians have the same genetic risk for AD as the general population. More than half of centenarians in population-based studies have cognitive impairment or dementia (46) and yet they still reach very old ages. This is in line with research that shows that about one-third of centenarians fit a “survivor” profile, meaning that they live 20 or more years with an age-associated chronic illness such as heart disease or cancer (47). Yet perhaps even more interesting are the centenarians who have pathological brain changes related to AD (i.e., buildup of amyloid and tau) and yet remain cognitively healthy, suggesting the presence of factors that promote resilience to AD (6,48). Various innate differences in brain structure and function as well as those incurred through life exposures (e.g., participation in cognitively stimulating, social, and physical activities) may underlie this and confer higher brain reserve (i.e., neurobiological differences of the brain that result in higher tolerance of AD pathology) or cognitive reserve (i.e., functional adaptation of cognitive processing in the presence of AD pathology (49)).

The analysis in this study reveals that centenarians are enriched of genetic variants protecting them from AD, beyond the genetic effects of *APOE*. Perhaps, an even more interesting finding in this study is that the genetic protection, as evidenced by the AD PPS, increased with increasing age. There was a non-significant increase in the AD PPS comparing nonagenarians and centenarians aged between 100 and 104 across NECS, LLFS, LGP, and 100-plus Study. A significant increase in the AD PPS was observed in a comparison between centenarians aged between 100 and 104 (mean AD PPS: 77.97) and semi-supercentenarians aged between 105 and 109 in the NECS (mean AD PPS: 78.58) and also in the TCS/JSS. Although not statistically significant, the same trend was observed in both LLFS and LGP. Lack of statistical significance in the LLFS and LGP could be attributed to reduced power as there were only 59 and 93 semi-supercentenarians, respectively, in the LLFS and LGP, while there were 516 semi-supercentenarians in the NECS and 577 semi-supercentenarians in the TCS/JSS. It is also worthwhile to point out that the supercentenarians (mean AD PPS: 79.14) had even higher AD PPS than semi-supercentenarians in the NECS, although this difference failed to reach statistical significance. Of note, the magnitudes of the difference between centenarians and semi-supercentenarians and between semi-supercentenarians and supercentenarians was 0.61 and 0.56, which show that differences are comparable in magnitude. Taken together, this analysis suggests that individuals who achieve the most extreme ages confer the greatest protection against AD. Genetic comparisons of these exceptional individuals to other groups further highlights the value of studying participants selected for extreme longevity.

The results from the NECS also showed that both the centenarians and offspring of centenarians had significantly higher AD PPS than the study controls, but there was no significant difference between centenarians and the offspring. Lack of replication in other cohorts may be due to participant recruitment differences and differences in the control group (i.e. spouses vs. offspring of parents with average survival vs. controls from other studies). Additionally, the NECS recruits the oldest participants and the offspring who were still alive to be recruited, who may have been the ones who had a lower genetic predisposition to AD. Increased genetic protection from AD among the offspring of centenarians is consistent with previous findings that the offspring has lower mortality (50) and longer disease-free survival (3,51) compared to the referent populations. Additionally, higher AD PPS among the offspring in our study may partially explain an extended cognitive health span in comparison to their peers that was observed in (52).

Based on the comprehensive dementia review in the LLFS, when AD PPS was correlated with the AD diagnosis in the LLFS, we observed AD PPS was not significantly predictive of AD (odds ratio=0.97, p=0.17), although the direction of effect was consistent with how the score was constructed. This may be due to censoring because most LLFS participants are still too young in this data set. For example, of the 1782 LLFS participants who underwent the dementia review, the vast majority of participants (n=1370) were in the offspring generation, and among these, only 47 were AD cases. Additional longitudinal follow-ups of these participants are warranted to find a more conclusive answer.

### Limitations

This study has some limitations. First, due to different genotype platforms and imputation panels, not all cohorts included in the study utilized the same set of SNPs in calculating the AD PPS. Accordingly, direct comparisons of the AD PPS across different cohorts were not possible in some cases. Second, with the exception of the NECS and TCS/JSS participants, there were limited sample sizes in the semi-supercentenarian and super-centenarian age groups. This may have resulted in unstable estimates of the AD PPS and low power in statistical comparisons. Third, examination of the range of the mean AD PPS across various groups in different cohorts reveals that the effect size of the genetic protection against AD is modest. For example, the increase in the AD PPS is in the magnitude of 1 or 2 additional protective alleles on average. Therefore, the results should be interpreted with caution. Lastly, when the BIMC test was administered to the centenarians in the NECS, those who became unable to complete the test due to cognitive impairment were not included in the current analysis, which may have biased the results.

### Conclusion

The current study presents clear evidence of genetic protection against dementia related to AD in centenarians compared to their respective study controls, despite their exceptional age which is a major risk factor for AD. Among the centenarians, it was also observed that the genetic protection increased with increasing age, particularly in the NECS and TCS/JSS. Perhaps, a future study should characterize the genetics of supercenternarians in the upper tail of the AD PPS distribution, who harbor more than a dozen protective alleles compared to their average peers.

## Supplementary Methods

### 100-plus Study / ADC / LASA

Cognitively healthy centenarians are included in the 100-plus Study if (i) they can provide official evidence for being aged 100 years or older, (ii) self-report to be cognitively healthy, which is confirmed by a proxy, (iii) consent to the donation of a blood sample, (iv) consent to (at least) two home visits from a researcher including an interview and neuropsychological test battery (10.1007/s10654-018-0451-3).

As healthy controls, we included (i) a sample of 1,773 Dutch older adults from the Longitudinal Aging Study of Amsterdam (LASA, 10.1007/s10654-016-0192-0), (ii) a sample of 1,524 older adults with subjective cognitive decline who visited the memory clinical of the Alzheimer Center Amsterdam and SCIENCe project, and were labeled cognitively normal after extensive examination (10.3233/JAD-170850), (iii) a sample of 62 healthy controls the Netherlands Brain Bank (10.1016/B978-0-444-63639-3.00001-3), (iv) a sample of 195 individuals from the twin study (10.1375/twin.13.3.231), and (v) a sample of 85 older adults from the 100-plus Study (partners of centenarians’ offspring, 10.1007/s10654-018-0451-3). Individuals with subjective cognitive decline were followed over time in the SCIENCe project, and only individuals who did not convert to mild cognitive impairment or dementia during follow-up were included in this study. The Medical Ethics Committee of the Amsterdam UMC approved all studies. All participants and/or their legal representatives provided written informed consent for participation in clinical and genetic studies.

